# Modeling Versatile and Dynamic Anaerobic Metabolism for PAOs/GAOs Competition Using Agent-based Model and Verification via Single Cell Raman Micro-spectroscopy

**DOI:** 10.1101/2020.11.18.387589

**Authors:** Guangyu Li, Nicholas B. Tooker, Dongqi Wang, Varun Srinivasan, James L. Barnard, Andrew Russell, Beverley Stinson, Jim McQuarrie, Peter Schauer, Adrienne Menniti, Erika Varga, Hélène Hauduc, Imre Takács, Charles Bott, Paul Dombrowski, Annalisa Onnis-Hayden, April Z. Gu

## Abstract

Side-stream enhanced biological phosphorus removal process (S2EBPR) has been demonstrated to improve performance stability and offer a suite of advantages compared to conventional EBPR design. Design and optimization of S2EBPR require modification of the current EBPR models that were not able to fully reflect the metabolic functions of and competition between the polyphosphate-accumulating organisms (PAOs) and glycogen-accumulating organisms (GAOs) under extended anaerobic conditions as in the S2EBPR conditions. In this study, we proposed and validated an improved model (iEBPR) for simulating PAO and GAO competition that incorporated heterogeneity and versatility in PAO sequential polymer usage, staged maintenance-decay, and glycolysis-TCA pathway shifts. The iEBPR model was first calibrated against bulk batch testing experiment data, being proved to perform better than the previous EBPR model for predicting the soluble orthoP, ammonia, biomass glycogen, and PHA temporal profiles in a batch starvation testing under prolonged anaerobic conditions. We further validated the model with another independent set of batch anaerobic batch testing data that included high-resolution single-cell and specific population level intracellular polymer measurements enabled by the single-cell Raman micro-spectroscopy technique. The model accurately predicted the temporal changes in the intracellular polymers at cellular and population levels within PAOs and GAOs, and further confirmed the proposed mechanism of sequential polymer utilization, and polymer availability-dependent and staged maintenance-decay in PAOs. These results indicate that under extended anaerobic phases as in S2EBPR, the PAOs may gain competitive advantages over GAOs due to the possession of multiple intracellular polymers and the adaptive switching of the anaerobic metabolic pathways that consequently lead to the later and slower decay in PAOs than GAOs. The iEBPR model can be applied to facilitate and optimize the design and operations of S2EBPR for more reliable nutrient removal and recovery from wastewater.

## INTRODUCTION

The enhanced biological phosphorus removal (EBPR) process has been recommended as a promising strategy to achieve sustainable wastewater P removal and facilitate P recovery (Gu et al., 2018). Current EBPR systems are designed to enrich a key functional group, namely the phosphate-accumulating organisms (PAOs) and *Ca. Accumulibacter phosphatis* is one of the most found PAOs in EBPR systems. Glycogen-accumulating organisms (GAOs) are often observed to coexist with PAOs, who possess a similar glycogen-based VFA-PHA metabolism without the polyphosphate (polyP) storage ability (Oehmen et al., 2006; Zeng et al., 2003) and are therefore concerned to be niche VFA (carbon source) competitors with PAOs consequently impacting EBPR performance. PAO-GAO competition could be kinetically affected by pH, temperature, substrate type, dissolved oxygen, solids retention time (SRT), hydraulic retention time (HRT), feeding strategy (Filipe et al., 2001a; Gu et al., 2008; Lopez-Vazquez et al., 2009; Majed and Gu, 2011; Neethling et al., 2006; Oehmen et al., 2005a, 2005b; Onnis-Hayden et al., 2020a; Rodrigo et al., 1996; Whang et al., 2007), etc. Although the proliferation of GAOs has been associated with EBPR performance deterioration in many cases (Gu et al., 2008; López-V et al., 2007; Oehmen et al., 2007), the coexistence of GAOs presence is not necessarily detrimental to EBPR performance as long as PAOs are kinetically favored (Gu et al., 2018; Nielsen et al., 2019; Onnis-Hayden et al., 2020b).

The performance stability of EBPR has been a concern hindering its wider applications. An emerging technology that has been demonstrated to successfully address the common stability challenge with EBPR is side-stream RAS and mixed liquor hydrolysis/fermentation-based side-stream EBPR (S2EBPR) (Barnard et al., 2010; Copp et al., 2012; Gu et al., 2018; Onnis-Hayden et al., 2020b; Stevens et al., 2015; Stroud and Martin, 2001; Vale et al., 2008; Vollertsen et al., 2006; Wang et al., 2019b). S2EBPR refers to modified EBPR configurations that include a diversion of RAS or anaerobic mixed liquor to a side-stream reactor, where simultaneous VFA production via sludge hydrolysis and fermentation and PAO activity-related P release and carbon uptake occur. Compared to conventional EBPR design, the S2EBPR offers a suite of advantages, including influent-carbon-independent VFA sources for PAO feast, eliminating the influences of fluctuating influent loads, more controllable lower-redox environments, more complex VFA compositions that likely favor the selection of PAOs over GAOs, flexible configurations and potential reduction of carbon footprint and denitrification enhancement by diverting more influent carbon to denitrifiers (Gu et al., 2018; Lanham et al., 2013; Mielczarek et al., 2013; Onnis-Hayden et al., 2020b; Srinivasan et al., 2021; Stokholm-Bjerregaard et al., 2017; Wang et al., 2019b).

While full-scale processes demonstrated the potential promises and advantages of S2EBPR (Barnard et al., 2010; Copp et al., 2012; Onnis-Hayden et al., 2020b; Stevens et al., 2015; Stroud and Martin, 2001; Vale et al., 2008; Vollertsen et al., 2006; Wang et al., 2019b), existing knowledge gaps in the fundamental understanding of the biochemical mechanisms and microbial ecology involved in S2EBPR hampers its wider application and implementation. Design and optimization of S2EBPR require adequate EBPR models that can capture the underlying key mechanisms involved in the S2EBPR such as the VFA production via hydrolysis and fermentation, and PAO and GAO competition in the extended anaerobic side-stream reactor. A few recent modeling efforts failed to predict either this competitive advantage or the performance superiority observed in S2EBPR systems compared to the conventional EBPR systems (Dunlap et al., 2016; Houweling et al., 2010). This suggested that there are still critical aspects in the current EBPR models that cannot reflect the metabolic functions of and competition between the PAOs and GAOs under the S2EBPR conditions, particularly the seemingly more pronounced differential metabolic versatility and dynamics among PAO and GAO subphenotypes under the prolonged anaerobic condition that involve cell maintenance, biomass decay and the associated versatility in the utilization of various intracellular polymers (Guedes da Silva et al., 2020; Lanham et al., 2014; G. Li et al., 2018; Santos et al., 2020).

First, PAO and GAO’s cell maintenance and cell decay are two processes that are highly relevant in S2EBPR, yet they are not fully understood or accurately modeled in previous EBPR models such as ASM (Henze et al., 2001). Cell maintenance is an observation-driven, hypothesized metabolic process. It was evidenced by continuous consumption of related intracellular polymers in PAO and GAO cells to produce ATP at a constant rate, throughout both short-term (i.e. ≤ 10 hrs) (Filipe et al., 2001a, 2001b; Houweling et al., 2010; Mino et al., 1998; Smolders et al., 1994a; Wentzel et al., 1989; Zeng et al., 2003) and long-term (≥5 days) (Vargas et al., 2013) anaerobic conditions while their known EBPR-related metabolisms are in idle. Previous models (e.g. ASM2 and ASM3+BioP (Henze et al., 2001)) often disregard the constant APT production rate but use a first-order polymer degradation to approximate, potentially causing deviated prediction of polymer consumption in cell maintenance. Evidence from previous literature (Carvalheira et al., 2014; Nielsen et al., 2016) and our recent studies (Gu et al., 2018; Wang et al., 2019b) suggested that the possession and ability of PAOs to utilize both polyP and glycogen offer them competitive advantages over GAOs that rely solely on glycogen, in terms of its survival time length, delayed decay, on-set time, and overall lower decay rates under extended anaerobic conditions as those in S2EBPR processes. This indicates that models such as Barker&Dold and UCTPHO+ (Barker and Dold, 1997; Hu et al., 2007), which do not account for the contribution of glycogen in PAO cell maintenance, can potentially underestimate PAOs’ competitive advantage. Recent models introduced a *sequential polymer usage* strategy to simulate cell maintenance (Lanham et al., 2014; G. Li et al., 2018; Santos et al., 2020) at constant ATP rates, and a *staged maintenance-decay* mechanism to approximate delayed cell decay when PAO and GAO have ample intracellular polymer for maintenance (G. Li et al., 2018; Santos et al., 2020). The previously observed lower-than-expected PAO and GAO decay rates (G. Li et al., 2018; Li et al., 2020; Santos et al., 2020) could potentially be explained by this maintenance survival hypothesis. However, two potential issues remain. First, the preference order of polyP and glycogen consumption is critical in the sequential polymer usage strategy, and both PAO subphenotypes may exist (Lopez et al., 2006; Lu et al., 2007). Conventional population-level models often assume one priority order (usually glycogen-prioritized) and ignore the other (Lanham et al., 2014; Santos et al., 2020), instead of considering both subphenotypes. Second, accurate simulation of cell decay among different PAO subphenotypes depends on the intracellular polymer’s availability at the cell level with the staged maintenance-decay strategy, which would be difficult to capture with conventional bulk measurements of aggregated polymer content in the biomass. Validating these proposed mechanisms requires both finer resolution measurements and simulation of the cellular polymer consumption heterogeneity among PAO subphenotypes.

Second, both overall microbial community and PAOs diversity in S2EBPR were found to be higher than conventional EBPRs (Onnis-Hayden et al., 2020b; Srinivasan et al., 2021). The recent discoveries pointed to highly varying and versatile anaerobic metabolic strategies among PAO subphenotypes (Guedes da Silva et al., 2020). (Santos et al., 2020) introduced the modeling of pathway switching for better simulating PAO metabolism. One of the most debated aspects related to the Accumulibacter’s anaerobic metabolism is the source of reducing power (NAD(P)H), which is required to reduce VFA into PHA monomers. Early studies suggest glycolysis and tricarboxylic acid (TCA) cycle to be possible sources, resulting in different PAO metabolic models per experimental evidence, including using sole glycolysis (referred to as Mino model) (Mino et al., 1987; Smolders et al., 1994a), sole TCA (referred to as Comeau-Wentzel model, C-W model) (Comeau et al., 1986; Maurer et al., 1997; Wentzel et al., 1989) or a mixture of the two in various ratios (Pereira et al., 1996; Pijuan et al., 2008; Yagci et al., 2003; Zhou et al., 2009). The switch between glycolysis and TCA-oriented metabolisms was hypothesized and implied to be triggered by intracellular polymer availability and depletion (Damir Brdjanovic et al., 1998; Guedes da Silva et al., 2020; Pijuan et al., 2008; Zhou et al., 2009).

In this study, to improve the EBPR model to better predict the S2EBPR processes, we proposed a revised EBPR model that better captures the PAO-GAO competition under extended anaerobic conditions, by incorporating staged maintenance-decay and glycolysis-TCA pathway shifts that are linked to the heterogeneity and versatility of PAO metabolism involving dynamic intercellular polymers usage. In addition, we employed an agent-based model to overcome the limitations in the current EBPR model that cannot capture the evidenced higher diversity of PAOs in S2EBPR than conventional EBPR (Srinivasan et al., 2021). The higher phylogenetic and phenotypic diversity of PAOs in the S2EBRP system compared to the conventional systems (Srinivasan et al., 2021; Wang et al., 2019b) likely resulted from more complex substrates and higher anaerobic SRT in the side-stream reactors and, they play a key role in the mechanisms underlying the PAO-GAO competition in S2EBPR. Agent-based modeling can capture the detailed PAO-GAO competition that involves the heterogeneity of subphenotypes (Y. Li et al., 2018; Majed et al., 2009; Majed and Gu, 2020, 2010) beyond those evaluated at aggregated and bulk estimates as in most previous studies (Guedes da Silva et al., 2020; Lopez-Vazquez et al., 2009). Furthermore and most importantly, we experimentally verified the proposed model with observations on cell-level, quantitative polymer dynamics by leveraging the power of single-cell Raman micro-spectroscopy technology (Bucci et al., 2012; Majed et al., 2009; Majed and Gu, 2011). To our knowledge, this was the first-time effort to verify the improved metabolic model using experimentally obtained single-cell intracellular polymer measurements from full-scale S2EBPR systems. The proposed model, referred to as iEBPR, outperformed the previous model in predicting the experimentally observed trends of intracellular polymer biomass content under anaerobic conditions and was proven to be more effective in simulating PAO and GAO maintenance behavior under those extended anaerobic conditions than conventional models. The proposed mechanism can be incorporated into future EBPR models to reveal the overall EBPR performance and PAO-GAO competitive dynamics more accurately as observed in S2EBPR systems.

## METHODOLOGY

### Agent-based EBPR Model Structure

An agent-based EBPR model (named iEBPR) was developed based on our previous iAlgae model developed by (Bucci et al., 2012). In this study, three population groups were included, namely PAOs, GAOs, and OHOs. OHOs stand for ordinary heterotrophic organisms that account for all non-PAO and non-GAO species. The same fundamental metabolic framework in iAlgae was used, including Accumulibacter-PAOs and OHOs taken from the International Water Association’s Activated Sludge Model 2 (ASM2) (Henze et al., 2001), and Competibacter-like GAOs coded for anaerobic VFA-uptake, PHA synthesis and aerobic biomass growth PHA-degradation and glycogen-accumulation (Filipe et al., 2001b; Houweling et al., 2010; Zeng et al., 2003). Note that iAlgae simplified VFAs to acetate and PHAs to PHB, which were kept in iEBPR and similar to (Santos et al., 2020). In addition, this study focused on PAO-GAO competition under extended anaerobic conditions; and denitrifiers, nitrifiers, and other anoxic-related organisms were not included. Similar to iAlgae, the inert organic matter was not tracked, and released matter from cell lysis would instantly transform into rbCOD (VFA) (Bucci et al., 2012) assuming the hydrolysis of inert organic matter is instant. The Gujer matrices of PAO, GAO, and OHO are shown in **Table S1-3**.

#### Representation of population phenotype heterogeneity

Each modeled species, i.e. PAO, GAO, or OHO was simulated in 10,000 independent agents known as the approach of agent-based modeling. Agents have independent polymer and biomass values and randomly assigned kinetic parameters to approximate metabolic heterogeneity among subphenotypic groups as in real activated sludge. Stoichiometric (yield) parameters were theoretically determined and were not randomized. Kinetic parameters, including rates and affinity parameters, were randomized by normal distributions, with their means representing the bulk-level aggregated values (for calibration to data, using literature values (Filipe et al., 2001b; Lopez-Vazquez et al., 2009; Zeng et al., 2003) or ASM2 defaults (Gujer, 1994)), and standard deviations for calibration. Invalid and exotic values (e.g. negative rates) from randomization were discarded and regenerated. In agent-based modeling, the number of agents is an important parameter to balance between simulating heterogeneity and computational time. We found that 1,000-10,000 agents per species were adequate considering this trade-off. In addition, all agents were updated synchronously at the end of each elapsed time step, known as the *discrete-event simulation* (Kurve et al., 2013).

### Polymer Availability-dependent Agent-level Energy Derivation for Cell Maintenance and Decay

Based on the fundamental metabolic framework from iAlgae, sequential polymer usage and staged maintenance-decay process were incorporated to replace the first-order decay mechanism. Unlike the population-level models in previous studies (Lanham et al., 2014; Santos et al., 2020), this calculation was performed at the agent level, allowing heterogenic cell maintenance and decay concerning the heterogeneity in polymer storage.

#### Cell maintenance with flexible and sequential polymer usage

PAO and GAO cell maintenance were simulated to fulfill a targeted ATP production rate, denoted as *m^ATP^*, expressed in mol-ATP/(C-mol biomass.hr). The equivalent polymer consumption rates to generate this ATP (in mg polyP-P/(mg biomass COD.day) and mg glycogen-COD/(mg biomass COD.day)) were calculated based on known stoichiometry and yield coefficients reported in the previous studies (Filipe et al., 2001b; Lu et al., 2007; Smolders et al., 1994a; Zeng et al., 2003). When the polymer(s) are ample, the actual maintenance is hypothesized to achieve the targeted rate (*m^ATP^*) without surplus; when polymer(s) are depleted, insufficient cell maintenance will occur and cause cell decay. This calculation is followed by both PAO and GAO biomass. In addition, as PAO has two polymers utilizable for cell maintenance, sequential polymer usage calculation is further incorporated to determine the individual ATP contribution from glycogen and polyP respectively. (Lanham et al., 2014) and (Santos et al., 2020) showed an example of this sequential usage calculation assuming using polyP prior to glycogen:

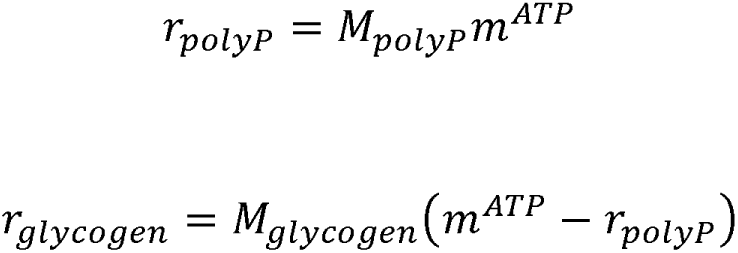

where: *r_polyP_* and *r_glycogen_* are the per polymer ATP contribution; *m^ATP^* is the target cell-maintenance rate; *M_polyP_* and *M_glycogen_* are Monod functions as rate modifiers based on per-polymer utilization affinity and availability. In this study, the other glycogen-preferring phenotype was also included

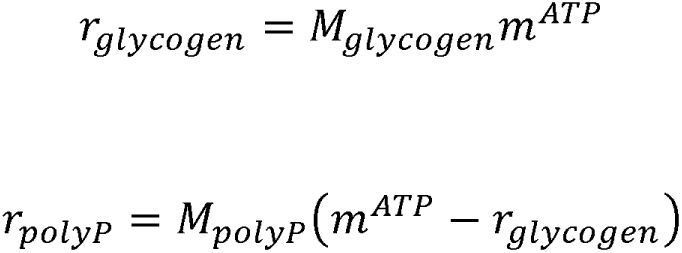

for enabling more flexibly simulating of the observed cases in various studies (Lopez et al., 2006; Lu et al., 2007; Vargas et al., 2013). The selection of polyP-glycogen priority is designed to be a calibratable parameter.

#### Staged and linked cell maintenance and decay

Previous EBPR models considered cell decay as an intrinsic process at a constant specific rate i.e., first-order decay, and as a process independent of cell maintenance. Several studies have shown evidence of accelerated biomass decay after depletion of their intracellular polymers. Since cell maintenance consumes polymers, this implies a linkage between biomass decay and cease of maintenance (Lu et al., 2007; Vargas et al., 2013). In our previous work (G. Li et al., 2018), we first introduced the process to link these two processes, hypothesizing that cells will aim to use the intracellular polymer(s)-derived energy for maintenance first to survive, before initiating decay when the energy can no longer satisfy the need for maintenance. Briefly, we treat the proportion of insufficient cell maintenance as the switch function to enable cell decay, making the PAO and GAO agents avoid decaying when their polymers are ample for cell maintenance, and start decaying when depleted:

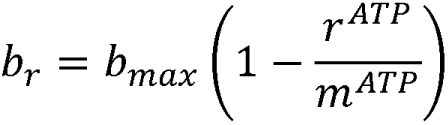

where: *_br_* is the actual specific decay rate; *b_max_* is the maximum specific decay rate. The formulation took into consideration of mathematical simplicity. The decayed biomass then proportionally regenerates as rbCOD (VFA) based on the empirical PAO/GAO biomass formula CH_1.93_O_0.53_N_0.2_ (Zeng et al., 2003). Although ammonia release was not direct tracked in the model, it could still be estimated using the above empirical biomass formula, which could be compared with experimental ammonia measurements used as an indicator of biomass decay (Lu et al., 2007). Linking cell maintenance to cell decay effectively models cell maintenance as an endogenous metabolic process that “precedes and inhibits” cell decay, making it a “survival strategy” for PAO and GAO species (Santos et al., 2020).

### Agent-level Glycolysis and TCA Cycle Pathway Switching

Based on our current knowledge, PAO’s glycolysis-oriented PHA synthesis will cease upon glycogen depletion, then the TCA-oriented PHA synthesis can continue as it doesn’t require glycogen (Guedes da Silva et al., 2020; Yagci et al., 2003; Zhou et al., 2009). The C-W model was chosen (Comeau et al., 1986; Wentzel et al., 1989) as the representative stoichiometry of TCA-oriented metabolism for its wide adoption in other EBPR models. The Mino model (Mino et al., 1998; Smolders et al., 1995) was used as the glycolysis-oriented stoichiometry. Note that both the Mino model and C-W model require the presence of polyP for PAO’s anaerobic PHA synthesis, making the same restriction in iEBPR. This implies that iEBPR does not allow GAO-like, polyP-independent metabolism in PAO cells, though it is still controversial in observation (Acevedo et al., 2012; Guedes da Silva et al., 2020). Furthermore, the glycolysis-oriented pathway was assumed to be prioritized due to its higher carbon efficiency than the TCA-oriented PHA synthesis (Guedes da Silva et al., 2020). To model the leveraged PHA synthesis rate, the following modifications are applied:

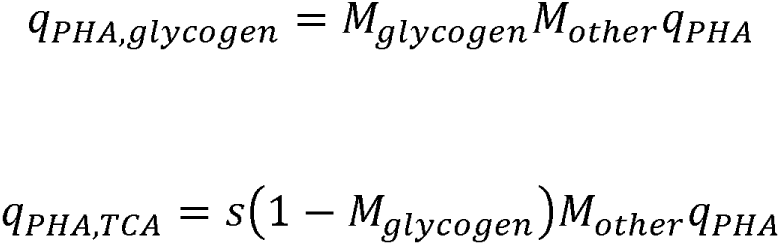

where *q_PHA,glycogen_* and *q_PHA,TCA_* are the adjusted PHA synthesis rates contributed by glycolysis-oriented and TCA-oriented metabolism respectively; *M_glycogen_* is the Monod function related to glycogen availability; *M_other_* is an aggregated term of other applicable rate modifiers used in ASM2; and *q_PHA_* is intrinsic PHA synthesis rate. The combined effect of these two PHA synthesis rates can simulate the employment of these two metabolism pathways in various degrees observed in previous studies (Comeau et al., 1986; Hesselmann et al., 2000; Lanham et al., 2013; Maurer et al., 1997; Mino et al., 1998; Pereira et al., 1996; Pijuan et al., 2008; Smolders et al., 1995; Wentzel et al., 1989; Yagci et al., 2003; Zhou et al., 2009). The binary parameter *s* (possible values are 0 and 1) is a phenotype indicator, for which 1 indicates an agent that can switch from glycolysis to TCA-oriented metabolism and 0 for being restricted to glycolysis-only. The relative abundance of agent subphenotypes can represent the relative abundance of PAO subphenotypes in real sludge, which enables the simulation of the coexistence and competition between these two subphenotypes. To our knowledge, this feature is exclusive to agent-based modeling and iEBPR is the first model to incorporate this into EBPR modeling.

### iEBPR Model Calibration

iEBPR was first calibrated with a set of data retrieved from a previous 8-day anaerobic starvation study in a lab-scale Accumulibacter-enriched (∼85% reported abundance) EBPR system (Lu et al., 2007). The initial contents of glycogen, PHA, and polyP were derived from the above literature using the empirical PAO biomass formula CH_1.93_O_0.53_N_0.2_ (Zeng et al., 2003). Gradient descent was used to automate the calibration of PAO and GAO anaerobic kinetic parameters. Briefly, the root mean square errors (RMSEs) of PHA, glycogen, orthoP, and MLVSS lysis (estimated by NH_4_ ^+^-N release) were calculated respectively; their geometric mean was used as the aggregated loss function (error measure). Geometric mean was used rather than arithmetic mean to arise the sensitivity of MLVSS lysis (NH_4_ ^+^-N release), which is two-magnitude smaller than orthoP. The calibration process takes a randomly generated parameter set as the start point and keeps adjusting the parameter values by small steps to reduce RMSE until no better parameter set can be found nearby (i.e. reached local minimum). Since local minima are sensitive to the initial parameter set, the above process was repeated 10,000 times with random initialization, and the best result was considered as the optimized parameter set. Other coefficients not subject to this calibration such as yields, were set the values from previous studies or the ASM2 model defaults (Lopez-Vazquez et al., 2009; Smolders et al., 1994b; Zeng et al., 2003). **Table S4-6** shows the final parameter set for PAOs and OHOs respectively, and unit conversions are shown in **Note S7**. The performance of the iEBPR model with calibrated parameter set was compared to conventional EBPR (calibrated separately).

### Cellular-level Experimental Observations via Single-cell Raman Micro-spectroscopy for Model Verification

Previous studies proposed single-cell Raman micro-spectroscopy (SCRS) to be a promising technology in estimating the glycogen, PHA, and polyP in individual cells, to further reveal the polymer distributions at both cellular and population levels (Y. Li et al., 2018; Majed et al., 2009; Majed and Gu, 2020, p. 20, 2010). This phenotypic survey data is comparable with the polymer distribution predicted by agent-based modeling (Bucci et al., 2012).

#### SCRS dataset acquisition

An SCRS dataset was first acquired individually for each sludge sample based on the method detailed in previous studies (Gu et al., 2018; Majed et al., 2009; Majed and Gu, 2011; Wang et al., 2019a). Briefly, 1 mL of MLSS was washed twice with 0.9% (w/v) NaCl solution and homogenized by passing in and out of a 26-gauge needle and syringe for at least 20 times to obtain uniform distribution of cells, as described previously. Then 6-8 drops of the disrupted sample were spread and dried on aluminum-coated slides (EMF Corp., Ithaca, US). After that, the slide was dipped into ice-cold Milli-Q water several times to remove salt particles and dried with filtered nitrogen gas. For each sample, Raman spectra for at least 80 single cells were acquired using a multiline confocal Raman spectrometer (LabRam HR Evolution, Horiba Jobin Yvon, Kyoto, Japan) configured with a 532 nm Nd:YAG laser and a 600 gr/mm grating. A 100× long working distance objective with a numerical aperture (NA) of 0.9 and a working distance of 0.21 mm was used to observe and collect Raman signals from single cells. The acquisition time for each spectrum was 20 seconds per cell and the laser power was set to 10%. Spectra were collected with scans from 400 cm^-1^ to 1800 cm^-1^.

#### SCRS data processing

Raman spectra processing and polymer relative abundance calculation were detailed in previous studies (Gu et al., 2018; Majed et al., 2009; Majed and Gu, 2011; Wang et al., 2019a). All Raman spectra were processed using cosmic spike removal, smoothing, background subtraction, and baseline correction using LabSpec 6 software (Horiba Jobin Yvon, Kyoto, Japan). Quality control was conducted by excluding the spectra showing unexpected signals (damaged) or low SNR, or lack of major characteristic peaks from bacterial components such as phenylalanine (∼1002 cm^-1^) and amide I (∼1657 cm^-1^). The candidate PAO and GAO populations were quantified based on the different combinations of intracellular polymeric inclusions, including poly-P (band at 690-700 cm^-1^ for P-O-P vibrations and band at 1168-1177 cm^-1^ for PO ^-^ stretching band), PHAs (bands at ∼434 cm^-1^,∼839 cm^-1^, and ∼1723 cm^-1^), and glycogen (bands at ∼480 cm^-1^,∼852 cm^-1^, and ∼938 cm^-1^), as described previously (Majed and Gu, 2020). The relative content of poly-P, PHAs, and glycogen in each candidate PAO and GAO cell were evaluated based on the intensity of the bands at 1168-1177 cm^-1^, ∼1723 cm^-1^, and ∼480 cm^-1^, respectively (normalized against the intensity of the amide I band). The polymer distribution in PAO and GAO species was then estimated based on the polymer relative abundances within PAO and GAO candidate cells collected in each sample respectively.

## RESULTS AND DISCUSSION

### iEBPR Model Calibrations against Anaerobic Starvation Batch Testing Results

iEBPR was calibrated against the temporal profile of orthophosphate and residual ammonia concentration and bulk measurements of intracellular glycogen and PHA using data extracted from (Lu et al., 2007, p. 200) (**Figure 1, Table 1**). The results showed that the sequential polymer usage and staged maintenance-decay as incorporated by the iEBPR performed quantitatively better than iAlgae/ASM2, in predicting the temporal profiles of orthoP release, cell lysis (NH_4_^+^-N release), glycogen, and PHA. Based on the experimental observations (**Figure 1**), the 8-day anaerobic starvation could be divided into 4 phases: (1) Glycogen degradation phase featuring fast glycogen reservoir depletion within the first day, accompanied by corresponding quick PHA formation and a relatively small amount of P release from polyP consumption; (2) Accelerated polyP consumption and P release, coupled with continuous glycogen degradation until its depletion. The majority of PHA accumulation (>90%) was done by the end of this phase in comparison to the final value observed at the end of this 8-day testing. The biomass (85% PAO) decay was also slight in this phase; (3) Deceleration of phosphate release indicated the slowing-down utilization of polyP, likely due to its near-depletion. The acceleration of NH_4_ ^+^-N release also indicated a transition of PAO endogenous metabolism and a starting of cell lysis. The acceleration then deceleration of orthoP release from Phase (1)-(3) resembled an S-shaped curve of orthoP concentration profile (**Figure 1**), indicating a transition in ATP source from glycogen to polyP based on glycogen availability. This transition was predicted by iEBPR but not by iAlgae/ASM2 which lacks the key mechanisms, such as sequential polymer usage.

**Figure 1.**
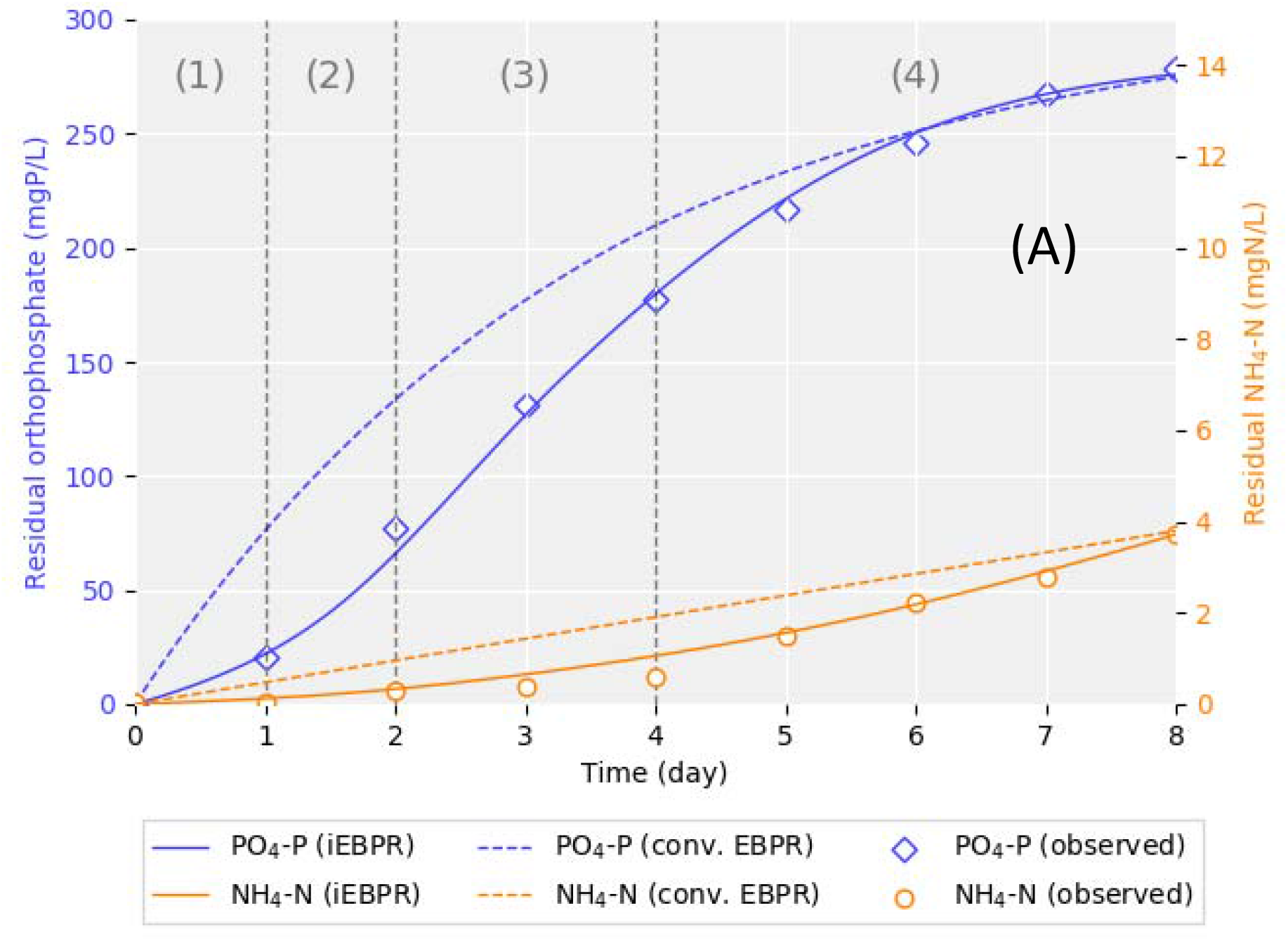
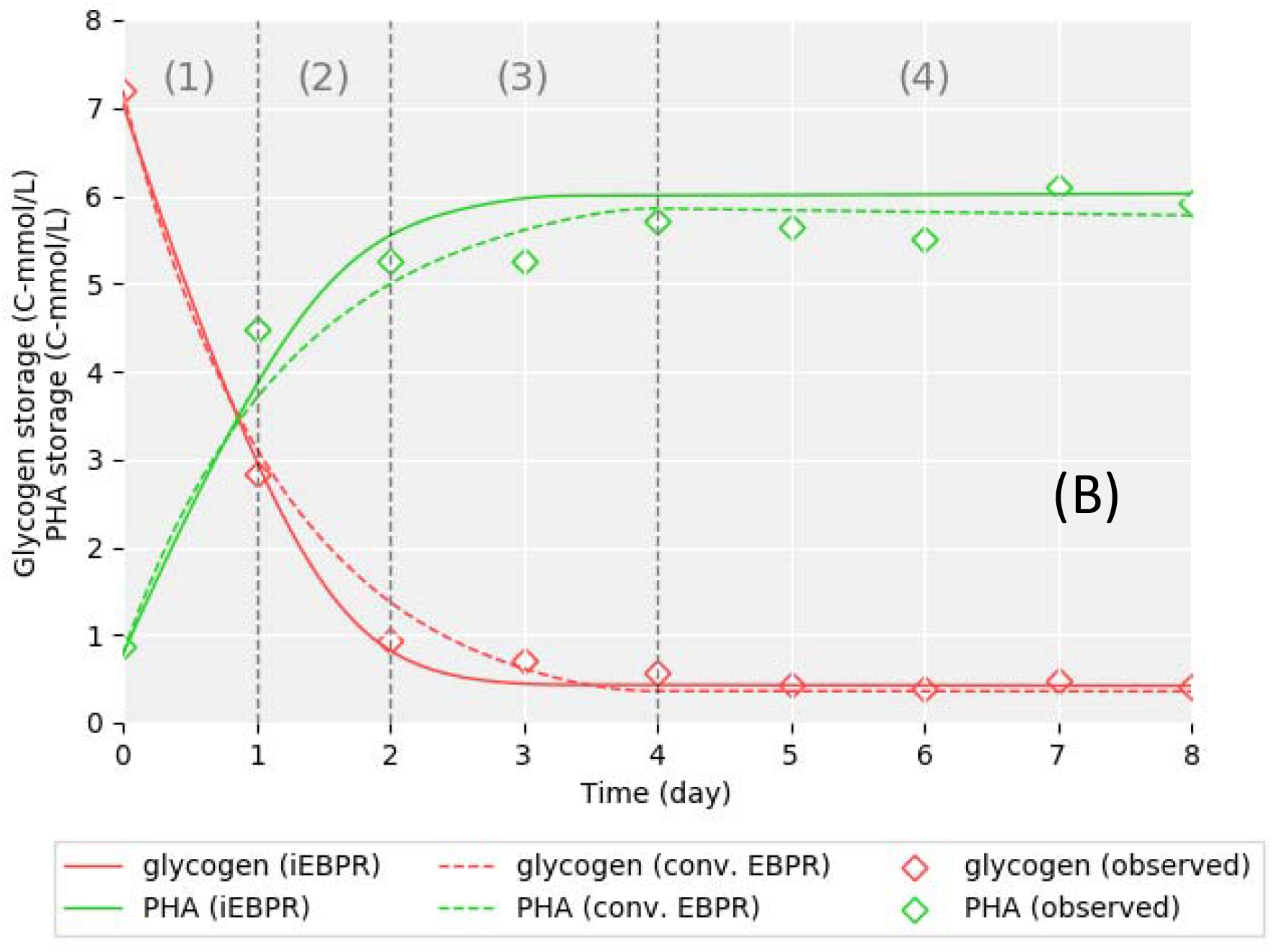
Model comparison between iEBPR and conventional EBPR in predicting the temporal profile of (A) orthoP and NH_4_^+^-N (cell decay indicator), and (B) intracellular glycogen and PHA, with experimental data acquired from an acetate-fed lab-scale EBPR system with approximately 85% Accumulibacter-PAOs during an 8-day anaerobic starvation batch test by (Lu et al., 2007). Time phases (1)-(4) indicate the four stages observed during the batch test: (1) rapid glycogen degradation; (2) transition stage from glycogen to polyP; (3) polyP utilization; and (4) polyP depletion stage with on-set of significant NH _4_^+^-N release (cell lysis).

**Table 1.** Modeling accuracy comparison in root mean square error (RMSE) between iEBPR and conventional EBPR (iAlgae/ASM2). Observation data was acquired from an 8-day anaerobic starvation test of an acetate-fed PAO-enriched (about 85%) lab-scale sludge by (Lu et al., 2007).

| Model | RMSE |  |  |  |
| --- | --- | --- | --- | --- |
|  | PO<sub>4</sub>-P | NH<sub>4</sub><sup>+</sup>-N | PHA | Glycogen |
|  | mgP/L | mgN/L | C-mol/L | C-mol/L |
| iEBPR* (this study) | 5.98 | 0.18 | 0.34 | 0.12 |
| iAlgae/ASM2(Bucci et al., 2012; Henze et al., 2001) | 33.0 | 1.08 | 0.34 | 0.19 |

\* Improved with staged maintenance-decay, sequential polymer usage in anaerobic cell maintenance, and glycolysis-TCA pathway switching.

**Figure 2** shows the detailed ATP productions suggesting a preference for glycogen priority over polyP for cell maintenance in this sludge. This was accompanied by the transition from glycogen-oriented to polyP-oriented cell maintenance, combined to achieve a cell maintenance rate at 2.1 × 10^−3^ mol ATP/(C-mol VSS·hr), which was comparable with values from other studies (**Table 1**). Calibration also suggested that only a very small portion of PAOs (< 1%) can switch to TCA-oriented PHA synthesis in this particular sludge, in agreement with the results found in the original experiment where no significant PHA formation was observed after glycogen depletion (Lu et al., 2007).

**Figure 2.**
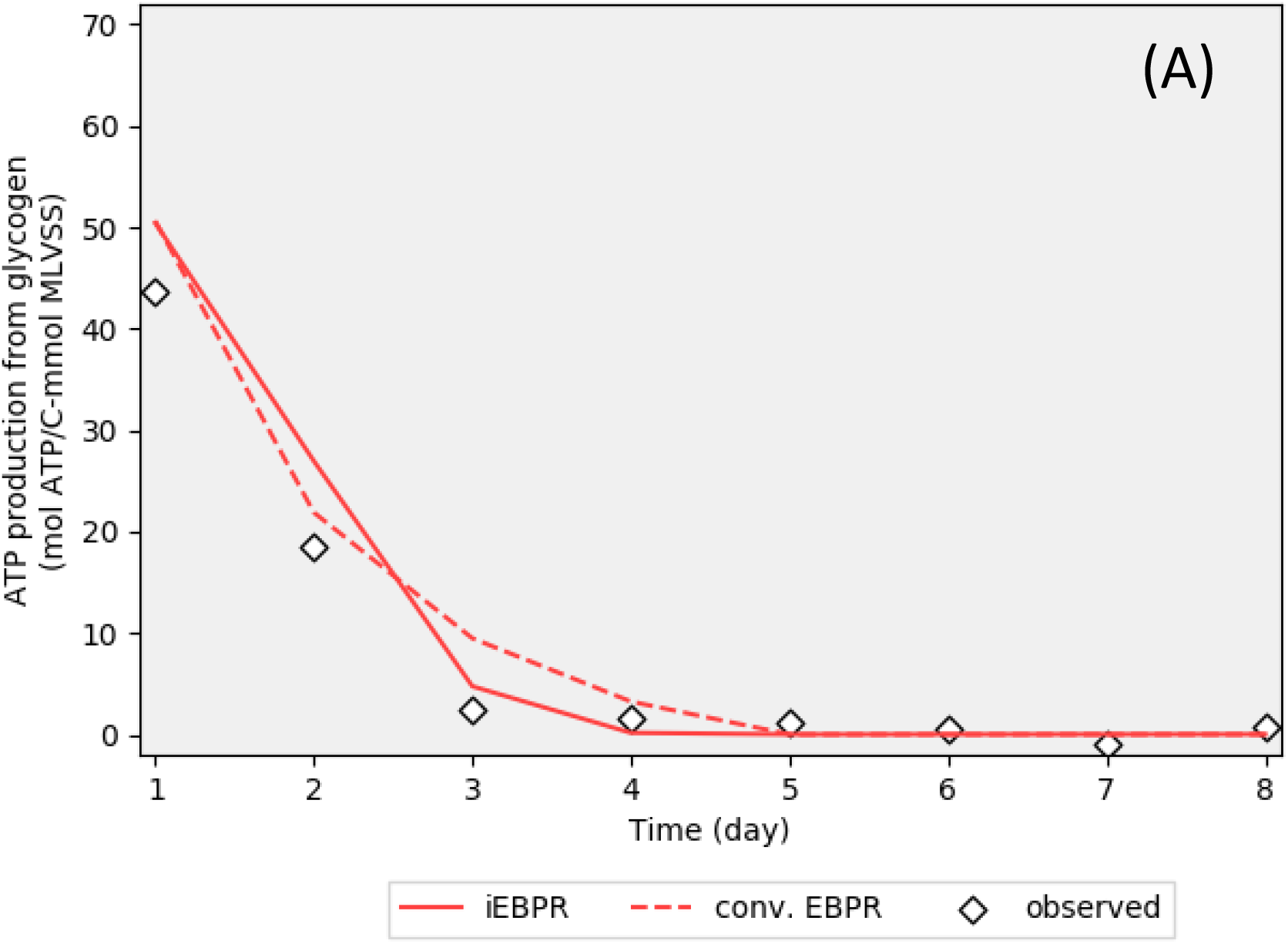
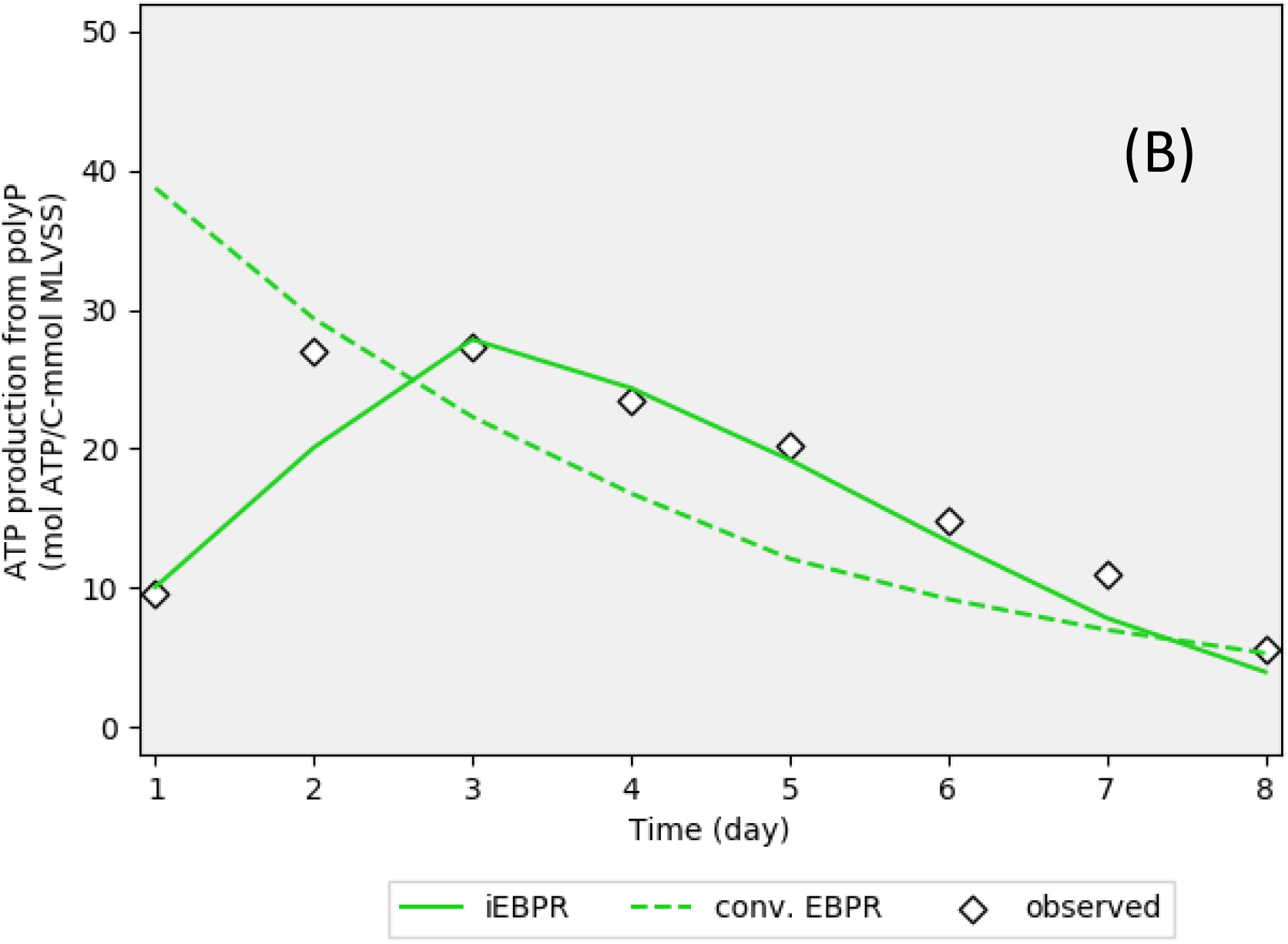
Comparison of model predictions and experimental observations of glycogen-contributed (A) and polyP-contributed (B) ATP during an 8-day anaerobic starvation testing with acetate-fed lab-scale EBPR system containing about 85% as Accumulibacter-PAOs (used as calibration dataset) by (Lu et al., 2007). The iEBPR developed in this study is designed with sequential anaerobic maintenance polymer usage, staged maintenance-decay, and glycolysis-TCA pathway switch; conventional EBPR (iAlgae/ASM2 (Bucci et al., 2012; Henze et al., 2001)) uses the same first-order decay.

#### Sequential polymer usage predicted PAO-GAO competition more accurately

Sequential polymer usage proposed a correction to the overestimation of anaerobic PAO intracellular polymer degradation. In comparison to the iAlgae which uses first-order decay kinetics, iEBPR estimated significantly less polyP degradation in the first three days during the batch testing (**Figure 1A**). The first-order polyP usage (used by iAlgae/ASM2) in cell maintenance failed to reflect the observed ATP production transition from glycogen to polyP-dependent, resulting in excessive polyP degradation when glycogen was still ample. The maximum polyP degradation overestimation was around 56 mgP/L in comparison to the observation data at the end of day 2 (**Figure 1**). This overestimation could significantly underestimate the PAO’s competitive advantage against GAO, particularly in the first 24 hrs. This could potentially explain the reason for conventional models like ASM2 to underestimate full-scale S2EBPR performances where prolonged anaerobic conditions are more relevant. A dedicated sequential polymer usage can therefore correct this overestimation and predict PAO anaerobic competition advantages more accurately.

#### Staged maintenance-decay predicted PAO-GAO competition more accurately

The staged maintenance-decay mechanism predicted more accurate cell decay kinetics, indicating an improved PAO anaerobic competitive advantage in comparison to first-order cell decay in iAlgae/ASM2 (**Figure 1&2, Table 1**) The observation data showed an acceleration in NH_4_ ^+^-N release at the end of day 4, which is adopted by other studies as a cell lysis indicator (Gujer, 1994; Lu et al., 2007). iEBPR could account for this acceleration using the hypothetical staged maintenance-decay mechanism, as shown in **Figure 1**. The calibrated PAO specific decay rate was 0.007/d, close to the reported value of 0.006/d from the literature. In comparison, iAlgae/ASM2 using conventional first-order decay cannot come up with a parameter value to comply with both short-term and long-term NH_4_ ^+^-N release predictions. For long-term (at the end of day 8) fit, a decay rate of 0.0028/d had to be used. Despite that this value was 53% below the literature value of 0.006/d, it caused a significant NH_4_ ^+^-N release overprediction of 1.91mg/L on day 4 (**Figure 1**) at the same time due to the assumption of always-occurring cell lysis implied by the first-order decay. In comparison, the relevant iEBPR prediction was 0.89mg/L, being much closer to the observed value of 0.60mg/L. Therefore, using more detailed cell decay kinetics, such as staged maintenance-decay could be crucial to improving the model performance and accuracy to simulate S2EBPR systems that have extended anaerobic incubation SRT.

### iEBPR Model Validation Case Study: Simulating PAO and GAO Competition under Extended Anaerobic Incubation Using Full-scale S2EBPR Sludge

#### Model prediction at the bulk level

The calibrated model was then used to simulate an independent anaerobic incubation batch testing similar to the conditions in the side-stream reactor in S2EBPR systems (Gu et al., 2018). The testing sludge was sampled from the side-stream fermentation reactor (in SSR configuration) of the South Cary Water Reclamation Facility (Apex, North Carolina) as described by (Gu et al., 2018) and (Onnis-Hayden et al., 2020a). It was estimated to contain 6.9% biovolume as Accumulibacter-PAOs and 1.1% as known GAOs by fluoresces *in situ* hybridization (FISH) (Gu et al., 2018). The model results were compared against the results from the batch tests that were conducted as a 72-hr anaerobic incubation without external VFA feeding, and the residual orthoP, residual VFA, MLSS glycogen, and PHA were monitored and measured at 0, 6, 12, 18, 24, 36 and 72 hours since the beginning of this incubation. The model was fitted to these temporal profiles with adjustment on the PAO/GAO/OHO composition, polymer contents, and the proportion of glycolysis-TCA switchable PAO phenotype. In addition, OHOs’ cell decay rate was also adjusted to the experimentally identified value. All other kinetic and stoichiometric parameters were set identical to the lab-scale sludge calibration results as described above in the previous section without further calibrations. GAO’s kinetic and stoichiometric parameters were set identical to PAO’s except for the yield ratio of PHA from VFA uptake, for which literature value derived from their glycogen-only metabolism was used (Lopez-Vazquez et al., 2009; Zeng et al., 2003).

**Figure 3** shows that the observed bulk measurements agreed well with the predicted temporal profiles of residual orthoP, residual VFA, total intracellular PHA, and glycogen during the anaerobic batch testing with S2EBPR sludge. The OHO decay rate was calculated to be 0.0076/d based on the VFA requirement to fit observed anaerobic metabolism, balanced by the stoichiometry of P release, glycogen consumption, and PHA formation. The model predicted a transitioning at around 45 hours from the high-rate active P release coupling a significant PHA synthesis phase to the second phase with a slower increase in both PHA and residual P. Calculated yield ratio of the active P release to PHA formation from 0-36 hours was 1.16 mol-P/C-mol PHA, which is comparable and within the range of 0.78-1.22 mol-P/C-mol PHA reported for S2EBPR sludge (Lanham et al., 2013) and it is higher than the typical range 0.36-0.77 mol-P/C-mol PHA for conventional A/O enriched sludge (Filipe et al., 2001a; Hu et al., 2007; Smolders et al., 1994a; Yagci et al., 2003). This high orthoP release is likely related to PAO metabolism as the orthoP release transition correlated with PHA formation, and potentially calls for adjustments and updates to the conventional PAO PHA synthesis stoichiometry for S2EBPR alike conditions. Overall, these bulk-level assessments demonstrated the general agreement of the model with mixture culture-level observations, though more detailed insights into the polymer fate and distribution at population and cellular levels that better reflect the competition between PAOs and GAOs could not be deduced.

**Figure 3.**
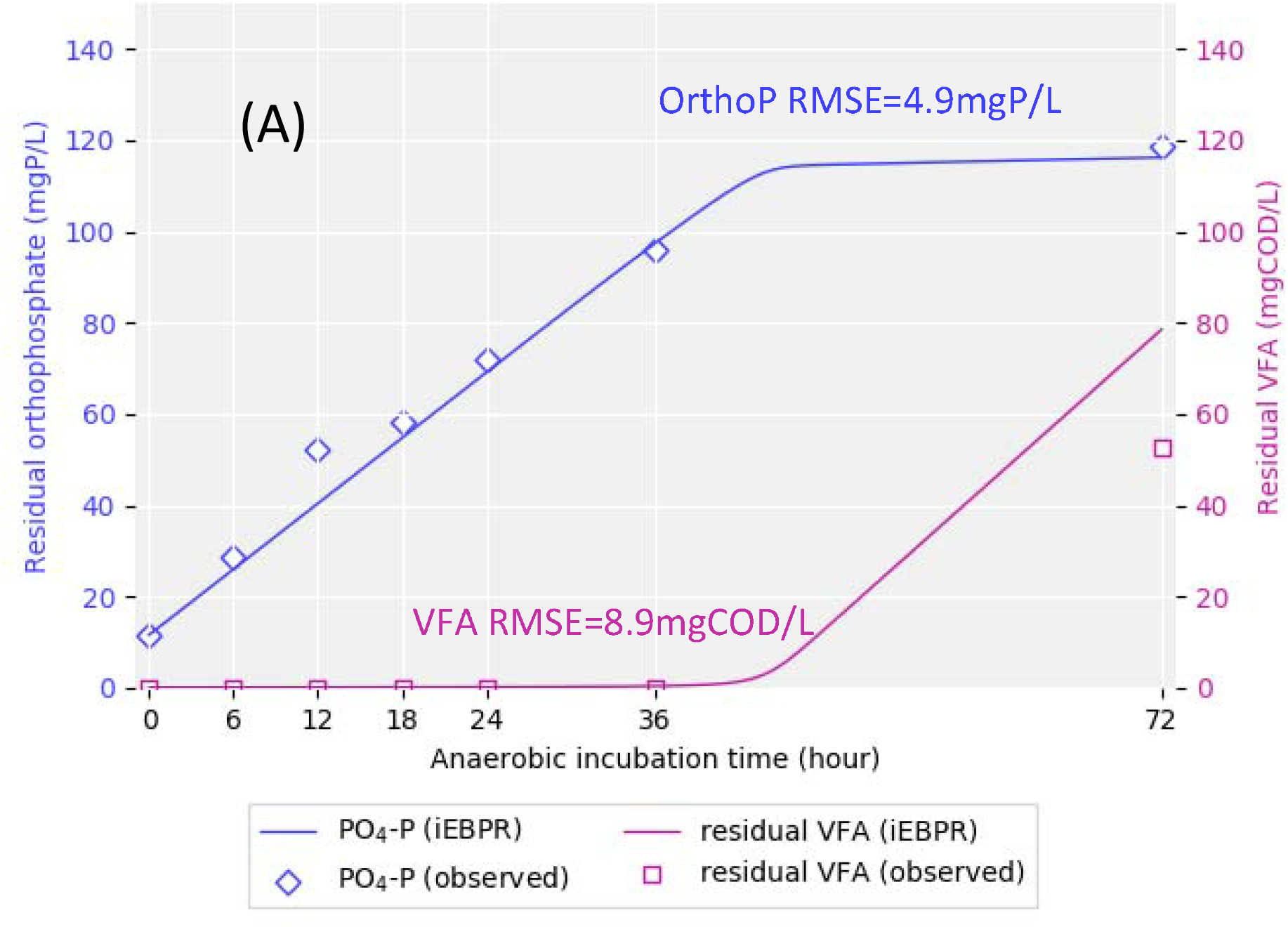
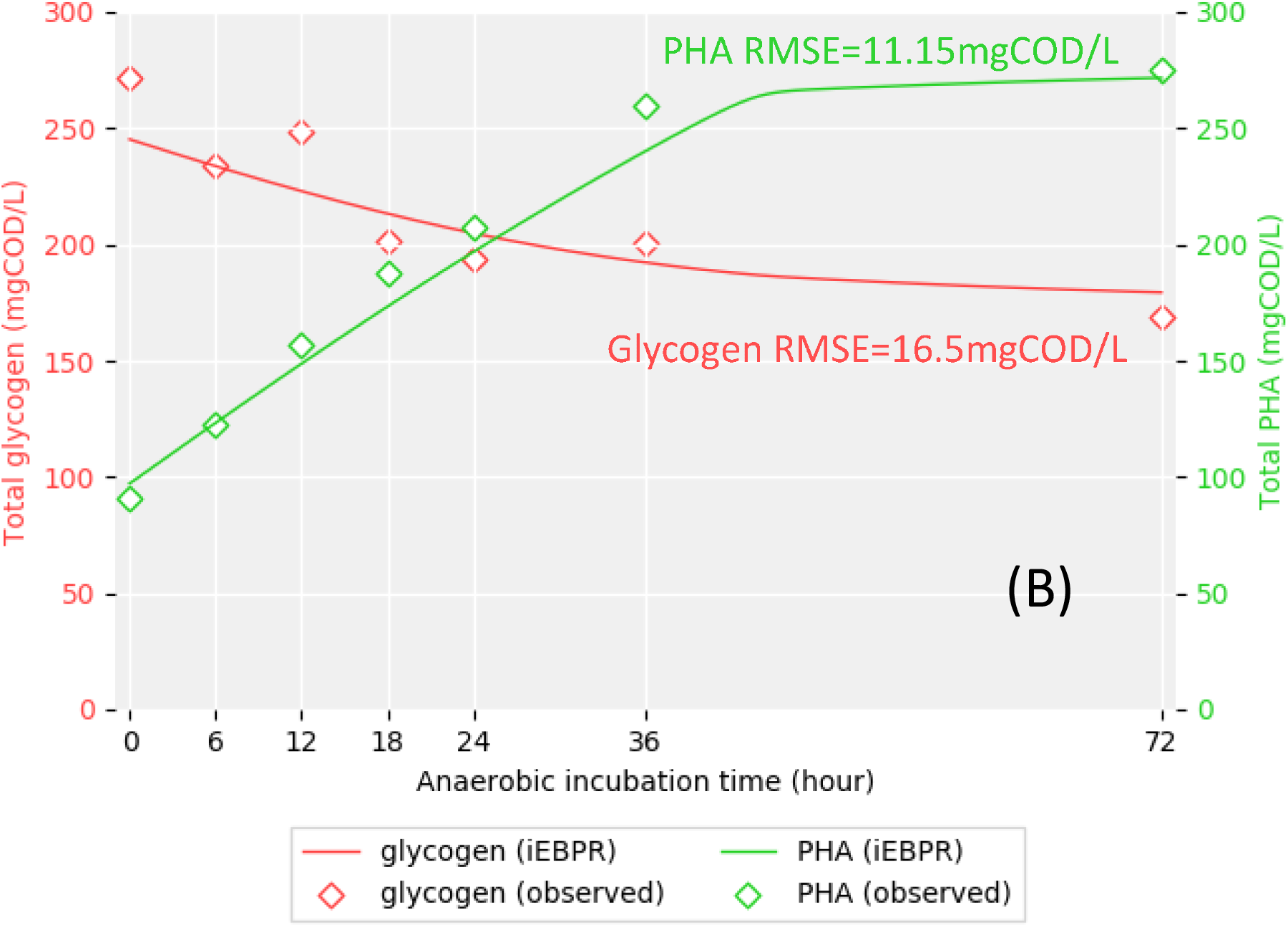
iEBPR simulation results of (A) residual orthoP and residual VFA, and (B) intracellular PHA and intracellular glycogen temporal profile in comparison to the experimental observations in a 72-hour anaerobic incubation batch testing. The tested activated sludge was sampled from a full-scale S2EBPR system (SSR configuration, South Cary WRF, Apex, NC), containing 6.9% as Accumulibacter-PAO, 1.1% as GAOs (estimated in biovolume by FISH). iEBPR incorporates three improved mechanisms: sequential polymer usage in cell maintenance, staged maintenance-decay, and PAO glycolysis-TCA pathway switch, all designed for S2EBPR-like extended anaerobic incubations.

#### Model prediction at the cell group level

The capability of inspecting intracellular polymer distribution at cell group resolution provided by the agent-based model enables an explicit way to further validate the proposed mechanisms related to polymer-dependent maintenance and staged maintenance-decay. SCRS was employed to measure and reveal the dynamics of intracellular polymers in PAOs and GAOs that ultimately dictate their competition, and they were compared against the agent-based model results. The initial distributions of PAOs’ and GAOs’ intracellular polymers, namely polyP and glycogen in PAOs and glycogen in GAOs, were fitted to the observed distribution following (Bucci et al., 2012) protocol. The model predicted polymer distributions were summarized from the agent polymer contents at 0, 6, 12, 24, 36, and 72 hours, and were compared with the SCRS observations at respective time points (Y. Li et al., 2018; Majed et al., 2009; Majed and Gu, 2010). The comparison between the model-predicted distributions against RCRS observations were shown in **Figure 4**. The accurate prediction of the EBPR-related intracellular polymer distribution changes indicates that the agent-based simulation in iEBPR can effectively track those polymer dynamics, emphasizing that our proposed maintenance-delay mechanism can be applied at the cell-group level and resolution.

**Figure 4.**
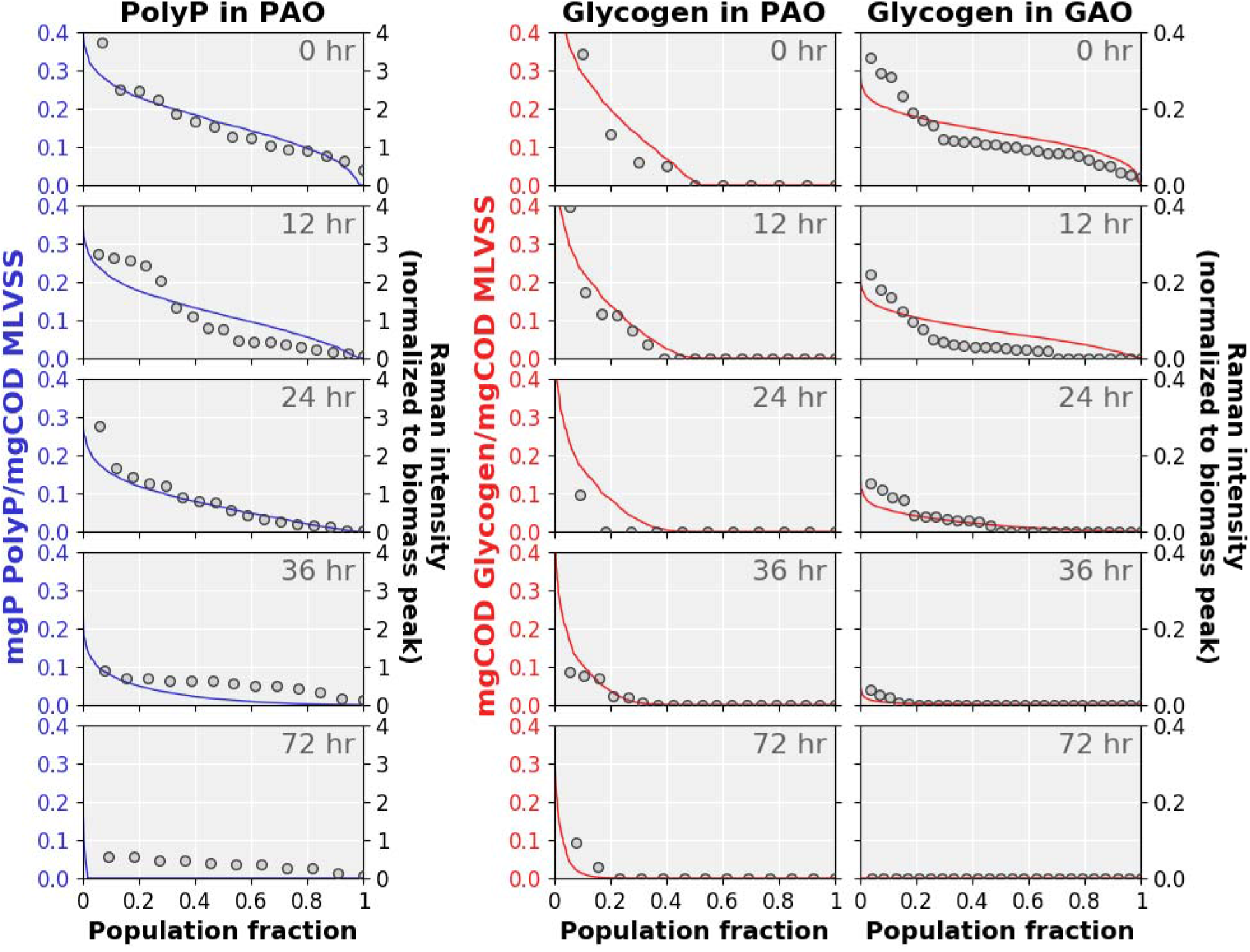
Comparison of model predicted polyP and glycogen dynamics in PAO and GAO cells with the observed dynamics acquired by Raman single-cell micro-spectroscopy (dots) during a 72-hr anaerobic incubation batch testing (this study). Tested sludge was sampled from a full-scale S2EBPR system (SSR configuration, South Cary WRF, Apex, NC)

#### PAO’s competitive advantage with polymer utilization versatility

It was hypothesized that the extended anaerobic condition and other factors in S2EBPR give PAOs more competitive advantages over GAOs (Carvalheira et al., 2014; Tooker et al., 2017; Wang et al., 2019b) Our results suggested that PAO’s more versatile polymer storage and metabolism were probable contributing factors to this advantage. **Figure 5** showed a glycogen depletion in GAO cells at around 36 hrs, with 97.2% of the initial storage already consumed. GAOs, therefore, transitioned into the decay status earlier than PAO because glycogen is the only energy source for their cell maintenance. In comparison, 33.9% of initial polyP and 25.3% of initial glycogen (19.3% and 30.3% as predicted by iEBPR respectively) were still available to PAOs for cell maintenance at 36 hours, with ∼92% and ∼16% of PAO cells still had detectable intracellular polyP and glycogen for cell maintenance as confirmed by the SCRS measurements. This indicated that the majority of PAO cells remained in the cell maintenance phase rather than transitioning into decay and lysis at 36 hrs. The model predicted ∼1.6% cumulative GAO biomass loss, in comparison to ∼1.2% PAO biomass decay. The possession and ability of PAOs to sequentially use multiple polymers enable its endurance to prolonged anaerobic conditions as shown in our study and previous studies (Carvalheira et al., 2014; Onnis-Hayden et al., 2020a; Vargas et al., 2013) likely contributed to PAOs’ competitive advantage over GAOs under S2EBPR conditions.

**Figure 5.**
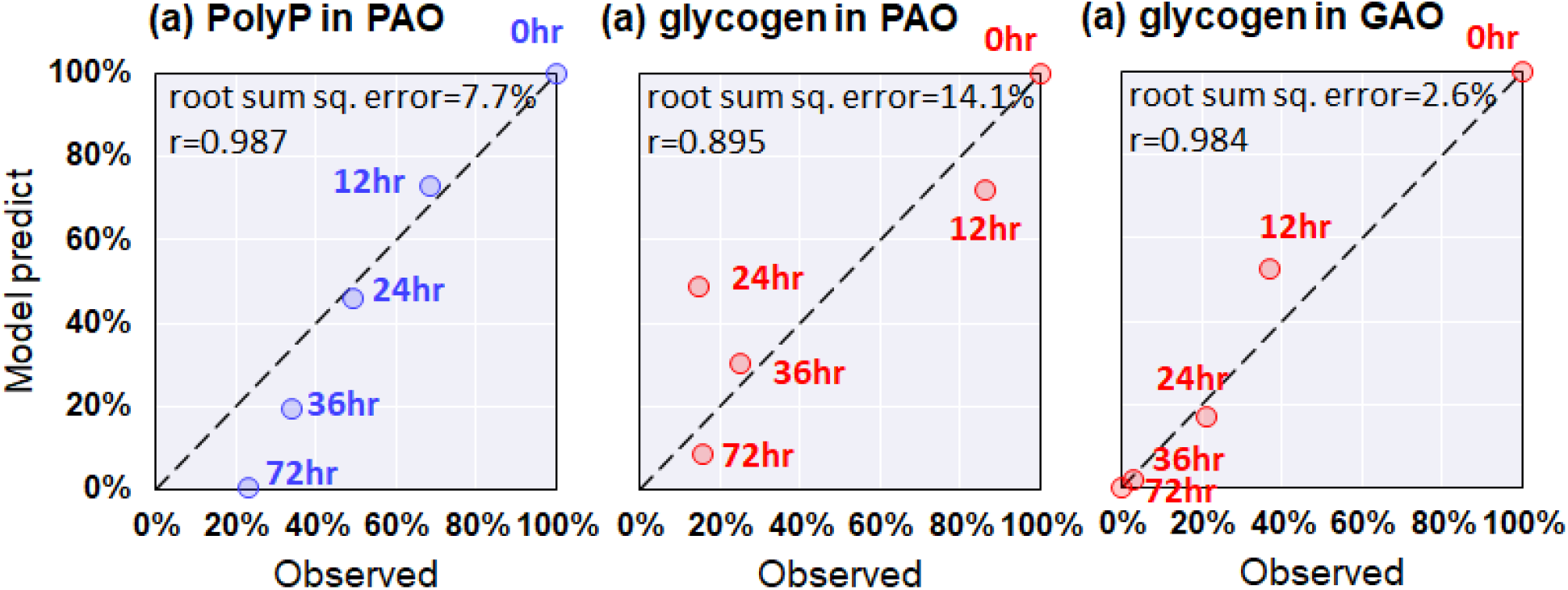
Comparison of model predicted polyP and glycogen dynamics in PAO and GAO cells with the observed dynamics acquired by Raman single-cell micro-spectroscopy during a 72-hr anaerobic incubation batch testing (this study). Polymer contents were normalized to their initial content at the beginning of this batch testing. Tested sludge was sampled from a full-scale S2EBPR system (SSR configuration, South Cary WRF, Apex, NC).

#### Modeling glycolysis-TCA pathway switch and PAO phenotype relative abundances

iEBPR provides a direct way to model, calibrate, and simulate the proportion of different PAO subphenotypes. Two PAO subphenotypes were modeled in iEBPR, (1) the glycolysis-TCA switchable PAO phenotype which switches to TCA-oriented metabolism and continues PHA synthesis when polyP is present, but glycogen is depleted, and (2) the glycolysis-only phenotype which the PHA synthesis will cease and transition into cell maintenance/decay when glycogen is depleted. The two subphenotypes coexist in iEBPR and by calibrating the abundance parameter of respective phenotype agents we can explicitly investigate and estimate the relative abundances of these two PAO subphenotypes within the activated sludge. Our result showed a dominance of glycolysis-TCA switchable phenotype, being calibrated to be 98.5%. The PHA accumulation would have been significantly less than the experiment observation when the cells were assumed to be strictly glycolysis-dependent, especially after the depletion of glycogen (**Figure 6**), in comparison to the calibrated fit when allowing glycolysis-TCA switching (**Figure 3**). This result agreed with previous observations that S2EBPR conditions potentially favor such PAO subphenotypes indicated by high anaerobic TCA cycle activity (Lanham et al., 2013). It is worth noting that iEBPR chose the C-W model to represent the TCA-oriented metabolic stoichiometry. In reality, it is a much more complex metabolic network composed of various shunts and branches (D Brdjanovic et al., 1998; He and McMahon, 2011; Hesselmann et al., 2000; Martin et al., 2006; Schuler and Jenkins, 2003; Wilmes et al., 2008; Yagci et al., 2003). A major challenge to modeling this complex metabolic network is to reliably determine the relative contribution of reducing power from each shunt and branch, which requires further elucidation by further experimental evidence.

**Figure 6.**
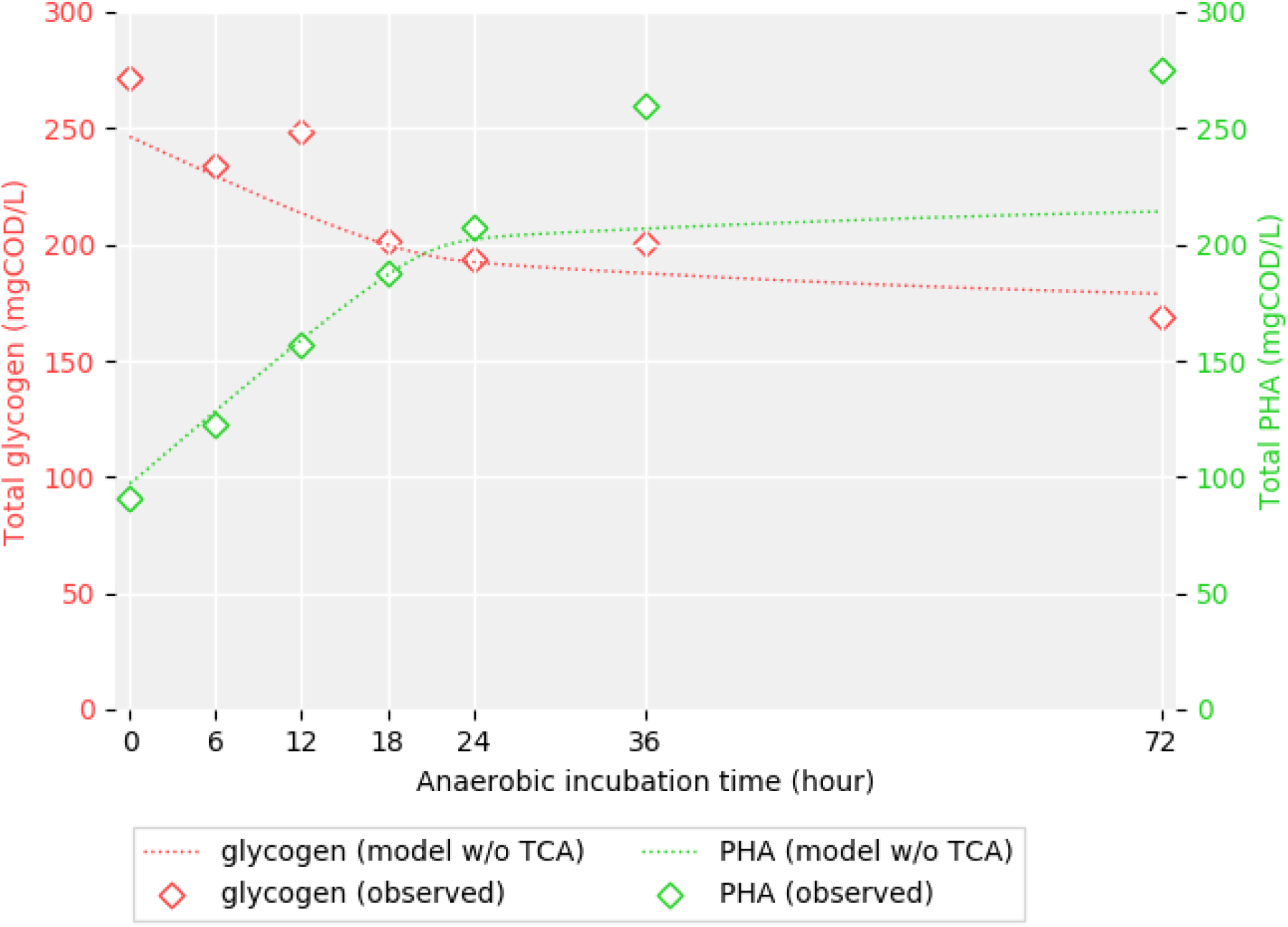
Modified iEBPR simulation restricting PAO cells to use glycolysis-only PHA synthesis resulted in significantly underpredicted PHA accumulation than observation. The tested activated sludge was sampled from a full-scale S2EBPR system (SSR configuration, South Cary WRF, Apex, NC), containing 6.9% as Accumulibacter-PAO, 1.1% as GAOs (estimated in biovolume by FISH). iEBPR incorporates three improved mechanisms: sequential polymer usage in cell maintenance, staged maintenance-decay, and PAO glycolysis-TCA pathway switch (disabled in this simulation), all designed for S2EBPR-like extended anaerobic incubations.

In addition, we acknowledge the limitations of iEBPR as an assessment tool to further resolve PAO phenotypes. First, non-Accumulibacter PAOs are not considered in iEBPR, such as the genus *Tetrasphaera* which does not accumulate PHA anaerobically (Fernando et al., 2019; Kristiansen et al., 2013). Therefore, we made the model framework expandable to incorporate new species in the future. Second, iEBPR does not account for “inert” polyP content that is experimentally observed (Barker and Dold, 1997). This could partially explain the remaining detectable polyP content persisted through the entire 72-hr incubation period (**Figure 4**). Third, iEBPR highly relies on measurements at the single-cell level resolution to accurately model the metabolic activity of an integrated microbial population. Breakthroughs in SCRS-based polymer quantification methods were still in demand to improve detection sensitivity and accuracy. Finally, competition on the substrates is the only modeled interaction between agents in iEBPR. Other interactions, either mutualistic or competitive, intraspecies or interspecies, are not considered in iEBPR but are common in the microbial ecology of activated sludge. More sophisticated SCRS datasets are required to reveal their effects which are beyond the scope of this study.

## CONCLUSION

iEBPR, an agent-based model that incorporates the sequential polymer usage, staged maintenance-decay processes, and glycolysis-TCA pathway shift, was developed, calibrated and validated using both bulk batch test experimental data and high-resolution cellular and population-level measurements enabled by SCRS.

- iEBPR improves the simulation accuracy on S2EBPR systems, where a longer anaerobic S2EBPR SRT may lead to more pronounced PAO-GAO anaerobic competition mechanisms that differ from those in play in conventional EBPR systems (e.g. A2O configuration).
- iEBPR revealed that under extended anaerobic conditions as in S2EBPR, the possession and versatile ability of PAOs to sequentially use different intracellular polymers (glycogen and polyP) as energy sources give them the competitive advantages over GAOs, as the results of longer survival with longer maintenance state and later on-set of cell decay than GAOs that can only utilize glycogen.
- iEBPR incorporates three newly proposed aspects, namely sequential polymer usage, staged maintenance-decay processes, and glycolysis-TCA pathway switch, which allows more accurate and higher-resolved simulation of the competition between PAO subphenotypes under anaerobic conditions.
- The combination of the agent-based model of coexistence of PAO subphenotypes of key populations and, SCRS-resolved intracellular polymer quantification allowed experimental verification of the proposed metabolic mechanisms at both population and cell group levels in activated sludge.
- iEBPR can be applied to facilitate and optimize the design and operations of S2EBPR for improved simulations of nutrient removal and recovery from wastewater.

Further studies are still in demand to improve iEBPR, including non-Accumulibacter PAOs, nitrogen-related metabolism and organisms, and modeling more diverse and comprehensive biological interactions between organisms.

## ACKNOWLEDGMENT

This work was supported by the Water Environment Research Foundation (Grant No. U1R13) and funds from Hampton Roads Sanitation District and Woodard & Curran, Inc. Special thanks to Clean Water Services and the South Cary Water Reclamation Facility operators and staff, and the entire WEF (previously WERF) S2EBPR Project Team for their generous support and assistance throughout the study.

