## Supplementary information for "Modeling Versatile and Dynamic Anaerobic Metabolism for PAOs/GAOs Competition Using Agent-based Model and Verification via Single Cell Raman Micro-spectroscopy"

**Table S1.** Gujer matrix of PAOs

|  | OP | VFA  (as acetate) | Biomass | PolyP | Glycogen | PHA | Kinetics |
| --- | --- | --- | --- | --- | --- | --- | --- |
| Ana. PHA synthesis, glycolysis | Y_P,release_ | -1 |  | -Y_P,release_ | 1-Y_HAc,PHA,gly_ | Y_HAc,PHA,gly_ | q_PHA_M_HAc_M_PP_M_gly_MI_PHA_ |
| Ana. PHA synthesis, TCA | Y_P,release_ | -1 |  | -Y_P,release_ |  | Y_HAc,PHA,TCA_ | q_PHA_M_HAc_M_PP_(1-M_gly_)MI_PHA_ |
| Ana. maintenance, polyP | 1 |  |  | -1 |  |  | m_PP_=m^ATP^M_PP_ (polyP-first)  m_PP_=m^ATP^(1-M_gly_)M_PP_ (glycogen-first) |
| Ana. maintenance, glycogen |  |  |  |  | -1 | 1 | m_gly_=m^ATP^M_gly_ (glycogen-first)  m_gly_=m^ATP^(1-M_PP_)M_gly_ (polyP-first) |
| Ana. biomass decay | i_BMP_ | 1 | -1 |  |  |  | b_ana_(1-(m_gly_+m_PP_)/m^ATP^) |
| Aer. glycogen synthesis |  |  |  |  | Y_PHA,gly_ | -1 | q_gly_M_PHA_MI_gly_ |
| Aer. polyP synthesis | -Y_PHA,PP_ |  |  | Y_PHA,PP_ |  | -1 | q_PP_M_PHA_M_P,storage_MI_PP_ |
| Aer. biomass growth | -i_BMP_ |  | Y_H_ |  |  | -1 | u_max_M_PHA_M_P,growth_ |
| Aer. biomass decay | i_BMP_ | 1 | -1 |  |  |  | b_aer_ |
| Aer. polyP decay | 1 |  |  | -1 |  |  | b_PP_ |
| Aer. glycogen decay |  |  |  |  | -1 |  | b_gly_ |
| Aer. PHA decay |  |  |  |  |  | -1 | b_PHA_ |

Notes:

* VFA, biomass, glycogen, and PHA are represented in mgCOD.

* M_<s>_ stands for Monod switch function of substrate S.

$$M_{S}=\frac{f_{S}-f_{S,min}}{K_{S}+f_{S}-f_{S,min}}$$

* MI_<s>_ stands for inhibitory Monod switch function of substrate S.

$${MI}_{S}=\frac{f_{S,max}-f_{S}}{{KI}_{S}+f_{S,max}-f_{S}}$$

* Parameters with identical names from PAO, GAO, and OHO are independent parameters.

* These statements apply to other Gujer matrices presented in this supplementary information document.

**Table S2.** Gujer matrix of GAOs

|  | OP | VFA  (as acetate) | Biomass | PolyP | Glycogen | PHA | Kinetics |
| --- | --- | --- | --- | --- | --- | --- | --- |
| Ana. PHA synthesis, glycolysis |  | -1 |  |  | 1-Y_HAc,PHA_ | Y_HAc,PHA_ | q_PHA_M_HAc_M_gly_MI_PHA_ |
| Ana. maintenance, glycogen |  |  |  |  | -1 | 1 | m_gly_=m^ATP^M_gly_ |
| Ana. biomass decay | i_BMP_ | 1 | -1 |  |  |  | b_ana_(1-m_gly_/m^ATP^) |
| Aer. glycogen synthesis |  |  |  |  | Y_PHA,gly_ | -1 | q_gly_M_PHA_MI_gly_ |
| Aer. biomass growth | -i_BMP_ |  | Y_H_ |  |  | -1 | u_max_M_PHA_M_P,growth_ |
| Aer. biomass decay | i_BMP_ | 1 | -1 |  |  |  | b_aer_ |
| Aer. glycogen decay |  |  |  |  | -1 |  | b_gly_ |
| Aer. PHA decay |  |  |  |  |  | -1 | b_PHA_ |

**Table S3.** Gujer matrix of OHOs

|  | OP | VFA  (as acetate) | Biomass | PolyP | Glycogen | PHA | Kinetics |
| --- | --- | --- | --- | --- | --- | --- | --- |
| Ana. biomass decay | i_BMP_ | 1 | -1 |  |  |  | b_ana_ |
| Aer. biomass growth | -i_BMP_ | -1 | Y_H_ |  |  |  | u_max_M_HAc_M_P,growth_ |
| Aer. biomass decay | i_BMP_ | 1 | -1 |  |  |  | b_aer_ |

**Table S4.** PAO parameter set

| Parameter | Definition | | Value | |
| --- | --- | --- | --- | --- |
|  |  |  | Calibration | Case study |
| f_gly_ | Glycogen content | [mgCOD/mgCOD MLVSS] | 0.114 | 0.036 |
| f_gly,max_ | Glycogen content, maximum quota | [mgCOD/mgCOD MLVSS] | 0.27 | 0.15 |
| f_gly,min_ | Glycogen content, minimum quota | [mgCOD/mgCOD MLVSS] | 0.007 | 0 |
| f_PHA_ | PHA content | [mgCOD/mgCOD MLVSS] | 0.014 | 0.012 |
| f_PHA,max_ | PHA content, maximum quota | [mgCOD/mgCOD MLVSS] | 0.4 | 0.4 |
| f_PHA,min_ | PHA content, minimum quota | [mgCOD/mgCOD MLVSS] | 0.01 | 0.01 |
| f_PP_ | PolyP content | [mgP/mgCOD MLVSS] | 0.142 | 0.143 |
| f_PP,max_ | PolyP content, maximum quota | [mgP/mgCOD MLVSS] | 0.28 | 0.28 |
| f_PP,min_ | PolyP content, minimum quota | [mgP/mgCOD MLVSS] | 0 | 0 |
| K_HAc_ | Half saturation constant, residual VFA (as HAc) | [mgCOD/L] | 3 | 3 |
| K_gly_ | Half saturation constant, glycogen utilization | [mgCOD/mgCOD MLVSS] | 0.03 | 0.03 |
| KI_gly_ | Inhibitory half saturation constant, glycogen synthesis | [mgCOD/mgCOD MLVSS] | 0.01 | 0.01 |
| K_PP_ | Half saturation constant, polyP utilization | [mgP/mgCOD MLVSS] | 0.044 | 0.044 |
| KI_PP_ | Inhibitory half saturation constant, polyP synthesis | [mgP/mgCOD MLVSS] | 0.001 | 0.001 |
| K_P_ | Half saturation constant, OP for biomass growth | [mgP/L] | 0.01 | 0.01 |
| K_P,storage_ | Half saturation constant, OP for polyP synthesis | [mgP/L] | 0.2 | 0.2 |
| K_PHA_ | Half saturation constant, PHA utilization | [mgCOD/mgCOD MLVSS] | 0.01 | 0.01 |
| Y_H_ | Biomass yield, heterotrophic | [mgCOD MLVSS/mgCOD] | 0.63 | 0.63 |
| Y_HAc,PHA,TCA_ | PHA yield per HAc uptake, TCA-oriented | [mgCOD/mgCOD HAc] | 1.00 | 1.00 |
| Y_HAc,PHA,gly_ | PHA yield per HAc uptake, glycolysis-oriented | [mgCOD/mgCOD HAc] | 1.50^[1]^ | 1.50^[1]^ |
| Y_PHA,gly_ | Glycogen yield per PHA oxidized | [mgCOD/mgCOD] | 0.75 | 0.75 |
| Y_PHA,PP_ | PolyP yield per PHA oxidized | [mgP/mgCOD] | 0.207 | 0.207 |
| Y_P,release_ | Active OP release with VFA uptake and PHA synthesis | [mgP/mgCOD] | 0.75 | 0.75^[2]^ |
| i_BMP_ | Biomass P content (exclude polyP) | [mgCOD/mgCOD MLVSS] | 0.02 | 0.02 |
| b_aer_ | Aerobic biomass decay rate | [d^-1^] | 0.032 | 0.032 |
| b_ana_ | Anaerobic biomass decay rate | [d^-1^] | 0.007 | 0.007 |
| b_gly_ | Aerobic glycogen decay rate | [d^-1^] | 0.0 | 0.0 |
| b_PHA_ | Aerobic PHA decay rate | [d^-1^] | 0.0 | 0.0 |
| b_PP_ | Aerobic polyP decay rate | [d^-1^] | 0.0 | 0.0 |
| m^ATP^ | Anaerobic maintenance requirement | [mol-ATP/C-mol MLVSS/d] | 0.05 | 0.05 |
| u_max_ | Maximum specific growth rate | [d^-1^] | 0.9 | 0.9 |
| q_gly_ | Maximum specific glycogen synthesis rate | [mgCOD/mgCOD MLVSS/d] | 1.8 | 1.8 |
| q_PHA_ | Maximum specific PHA synthesis rate | [mgCOD/mgCOD MLVSS/d] | 3.0 | 3.0 |
| q_PP_ | Maximum specific polyP synthesis rate | [mgP/mgCOD MLVSS/d] | 1.5 | 1.5 |
| p_TCA-switch_ | Fraction of agents with glycolysis-TCA pathway switching | dimensionless | 0.003 | 0.985 |

Notes:

[1] This is the overall yield ratio including PHA synthesized from VFA (HAc) and transformed products from glycogen.

[2] This value was reported by Lopez-Vazquez et al., (2009) for 1:1 HAc:HPr at pH=7.0.

**Table S5.** GAO parameter set

| Parameter | Definition | | Value | |
| --- | --- | --- | --- | --- |
|  |  |  | Calibration | Case study |
| f_gly_ | Glycogen content | [mgCOD/mgCOD MLVSS] | 0.097 | 0.04 |
| f_gly,max_ | Glycogen content, maximum quota | [mgCOD/mgCOD MLVSS] | 0.3 | 0.3 |
| f_gly,min_ | Glycogen content, minimum quota | [mgCOD/mgCOD MLVSS] | 0.005 | 0 |
| f_PHA_ | PHA content | [mgCOD/mgCOD MLVSS] | 0.012 | 0.0133 |
| f_PHA,max_ | PHA content, maximum quota | [mgCOD/mgCOD MLVSS] | 0.3 | 0.3 |
| f_PHA,min_ | PHA content, minimum quota | [mgCOD/mgCOD MLVSS] | 0.01 | 0.01 |
| K_HAc_ | Half saturation constant, residual VFA (as HAc) | [mgCOD/L] | 3 | 3 |
| K_gly_ | Half saturation constant, glycogen utilization | [mgCOD/mgCOD MLVSS] | 0.01 | 0.01 |
| KI_gly_ | Inhibitory half saturation constant, glycogen synthesis | [mgCOD/mgCOD MLVSS] | 0.02 | 0.02 |
| K_P_ | Half saturation constant, OP for biomass growth | [mgP/L] | 0.01 | 0.01 |
| K_PHA_ | Half saturation constant, PHA utilization | [mgCOD/mgCOD MLVSS] | 0.01 | 0.01 |
| Y_H_ | Biomass yield, heterotrophic | [mgCOD MLVSS/mgCOD] | 0.63 | 0.63 |
| Y_HAc,PHA_ | PHA yield per HAc uptake, TCA-oriented | [mgCOD/mgCOD HAc] | 2.21^[1]^ | 2.21^[1]^ |
| Y_PHA,gly_ | Glycogen yield per PHA oxidized | [mgCOD/mgCOD] | 0.75 | 0.75 |
| i_BMP_ | Biomass P content (exclude polyP) | [mgCOD/mgCOD MLVSS] | 0.02 | 0.02 |
| b_aer_ | Aerobic biomass decay rate | [d^-1^] | 0.032 | 0.032 |
| b_ana_ | Anaerobic biomass decay rate | [d^-1^] | 0.006^[2]^ | 0.006 |
| b_gly_ | Aerobic glycogen decay rate | [d^-1^] | 0.0 | 0.0 |
| b_PHA_ | Aerobic PHA decay rate | [d^-1^] | 0.0 | 0.0 |
| m^ATP^ | Anaerobic maintenance requirement | [mol-ATP/C-mol MLVSS/d] | 0.079 | 0.079 |
| u_max_ | Maximum specific growth rate | [d^-1^] | 0.9 | 0.9 |
| q_gly_ | Maximum specific glycogen synthesis rate | [mgCOD/mgCOD MLVSS/d] | 3 | 3 |
| q_PHA_ | Maximum specific PHA synthesis rate | [mgCOD/mgCOD MLVSS/d] | 3.5 | 3.5 |

Notes:

[1] This is the overall yield ratio including PHA synthesized from VFA (HAc) and transformed products from glycogen.

[2] This value was hypothesized to be the same as PAOs due to low abundance.

**Table S6.** OHO parameter set

| Parameter | Definition | | Value | |
| --- | --- | --- | --- | --- |
|  |  |  | Calibration | Case study |
| K_HAc_ | Half saturation constant, residual VFA (as HAc) | [mgCOD/L] | 4 | 4 |
| K_P_ | Half saturation constant, OP for biomass growth | [mgP/L] | 0.01 | 0.01 |
| Y_H_ | Biomass yield, heterotrophic | [mgCOD MLVSS/mgCOD] | 0.63 | 0.63 |
| i_BMP_ | Biomass P content (exclude polyP) | [mgCOD/mgCOD MLVSS] | 0.02 | 0.02 |
| b_aer_ | Aerobic biomass decay rate | [d^-1^] | 0.2 | 0.2 |
| b_ana_ | Anaerobic biomass decay rate | [d^-1^] | 0.006^[1]^ | 0.007 |
| u_max_ | Maximum specific growth rate | [d^-1^] | 6.0 | 6.0 |

Notes:

[1] This value was hypothesized to be the same as PAOs due to low abundance.

**Notes S7.** Unit conversion of maintenance requirements

**(1) Maintenance by polyP cleavage.**

Smolders et al., (1994) suggested that the P-mol ATP yield is approximately equal to P-mol polyP cleaved, i.e.

$$Y_{PP,ATP}=\frac{1 mol ATP}{1 mol PP-P}.$$

Therefore,

$$\frac{1 mol ATP}{1 C-mol VSS}=\frac{\frac{1 mol PP-P}{1 mol ATP}\times\frac{31 gP}{1 mol P}}{\frac{34.16 gCOD MLVSS}{1 C-mol VSS}}=\frac{0.907 gPP-P}{1 gCOD VSS}.$$

**(2) Maintenance by glycolysis via Embden-Meyerhof-Parnas (EMP) pathway.**

Filipe et al., (2001) and Lu et al., (2007) suggested below EMP pathway stoichiometry for pure ATP production was applicable for both Accumulibacter-PAOs and Competibacter-GAOs:

$$Y_{gly,ATP}^{ANA}=\frac{\frac{2}{3}mol ATP}{6 C-mol gly.}.$$

Therefore,

$$\frac{1 mol ATP}{1 C-mol VSS}=\frac{\frac{6 C-mol gly.}{2/3 mol ATP}\times\frac{192 gCOD gly.}{6 C-mol gly.}}{\frac{34.16 gCOD MLVSS}{1 C-mol VSS}}=\frac{8.43 gCOD gly.}{1 gCOD VSS}.$$
